# Integrating neurophysiological insights into effective bird deterrence using flickering light

**DOI:** 10.1101/2024.06.20.599983

**Authors:** Takeshi Hondaa, Hiroki Tominaga, Akio Shimizu

## Abstract

There are few effective methods to mitigate the economic or health-related disadvantages caused by birds. Traditional countermeasures employing sound and light have been utilized for mitigating crop damage, but their efficacy is insufficient, and human-avian conflicts persist. This study explores a fundamentally different approach to resolving these conflicts. Flashing lights that alternate between red and blue are known to stimulate the human brain and can potentially cause photosensitive epilepsy in one in 100,000 people, though very rarely. A 15 Hz flashing stimulus is known to elicit a significant response in humans; however, the optimal flashing pattern for birds remained unclear. We investigated the effect of different flickering patterns, specifically 12.5, 15, and 20 Hz, on crows when illuminated from a distance within 200 meters. The flashlight used was a long-range model and the Light Emitting Diodes (LEDs) consumed about 9 watts of power. The power was supplied by 21700 type lithium-ion batteries. Using a long-range flashlight during daylight, we determined that a 15 Hz flashing stimulus was most effective. This finding suggests that the most intense stimulus identified in human physiology can be equally effective when applied to birds. Survival analysis estimated that by projecting this pattern from a distance of 100 meters, crows fled within 8.1 seconds. Unlike traditional bird damage control techniques that rely on neophobia, this study utilizes physiological aversion. In this regard, our approach is fundamentally different from traditional techniques. The method of using flashing light to stimulate the brain, rather than the eyes, is based on insights from human medical and physiological studies. These findings elucidate the principle of a novel aversive stimulus using flashing light, which holds potential for widespread application in mitigating bird-related issues.

## 1. Introduction

The interactions between birds and humans have historically spanned a wide spectrum, resulting in a complex array of conflicts. These include economic impacts in agriculture, where birds damage cereals, fruits, and vegetables (Conover, 2001), as well as public health risks associated with avian influenza transmission (Lang et al., 2008). Technology to resolve these issues include isolation and deterrence. Isolation involves using physical barriers like nets to prevent birds from entering (Fuller-Perrine and Tobin, 1993), while deterrence aims to keep birds away from unwanted areas (Blackwell et al., 2002; Elbers and Gonzales, 2021; Savereno et al., 1973). For isolation, bird nets are used, but covering vast agricultural lands with nets poses problems in terms of installation effort and cost, and it is not widely adopted (Fuller-Perrine and Tobin, 1993). Various technologies have been developed for deterrence, but maintaining long-term deterrence is challenging. For auditory deterrence, technologies such as propane cannons (Conover, 1984; Drake and Villano, 2005), guns (Heim et al., 2022; Månsson, 2017), and distress calls (Bomford and O ‘brien, 1990; Johnson et al., 1985) have been used; however, their effectiveness is either insufficient or significantly decreases over a short period. In summary, existing technologies are insufficient to reduce the damage caused by birds.

Aversive conditioning has been shown to be effective in guiding animals to avoid certain spaces (i.e., deterrence) (Avcu et al., 2014). However, to effectively induce aversion, the stimuli must involve real discomfort rather than simply being novel and unfamiliar. In birds, aversive conditioning can be induced using thiram as a chemical repellent (Clark, 1998; Kennedy and Connery, 2008). Conditioning is possible because animals that consume food containing this substance experience nausea. In wildlife management, electric stimuli such as electric fences maintain their deterrent effect as wildlife learn to associate the discomfort of electric shock with the fence. However, visual and auditory stimuli used in bird control often fail to provide sufficient discomfort, relying instead on their novelty (neophobia). For example, loud noises like those from a propane cannon may temporarily increase vigilance but considering that birds can become unresponsive to the sound of aircraft jet engines and cause bird strikes, it is not difficult to predict that loud noises will not have a long-term effect (Washburn et al., 2006). Thus, existing deterrence technologies face difficulty in effectively eliminating uncertainty.

Recent physiological studies have shown that light stimuli can induce discomfort in humans. The symptoms caused by this discomfort stimulus are known in medical science targeting humans as flicker vertigo or photosensitive epilepsy (Angus, 1995). Red flickering light is found to be more discomforting than white in monochromatic cases (Parra et al., 2007; Takahashi and Tsukahara, 1992). Furthermore, a combination of red and blue light is more stimulating than monochromatic light in dichromatic cases (Da Silva and Leal, 2017; Harding, 1998; Wilkins et al., 2010). This light stimulus acts as a physiological agent targeting the brain, rather than the eyes (Angus, 1995; Rash, 2004). While photosensitive epilepsy is a rare condition in humans, affecting one in 100,000 individuals (Da Silva and Leal, 2017), flicker vertigo or photosensitivity to flickering light is a well-known physiological response (Angus, 1995). Until recently, there has been limited knowledge on how flickering light affects birds, with some studies conducted on a special breed of chicken (Fayoumi) known for its high proportion of photosensitive epilepsy patients (Batini et al., 2004; Nunoya et al., 1983). However, to our knowledge, no research has been conducted to determine whether flickering light can act as an aversive stimulus for birds, similar to its effects on humans. If flickering light can induce physiological aversion in birds, it may be a viable technique for bird management.

In this study, we aim to explore the most effective flickering light stimulus for use as a damage control technology by varying the flickering frequency and pattern. If the effects of flickering light on birds and humans are the same, a 15 Hz flicker with alternating red and blue light should induce the most effective response. Identifying the most effective flickering pattern is essential for utilizing flickering light as a bird behavior control technology.

## 2. Methods

### 2.1. Study area

In this study, we assessed the impact of flickering stimuli on carrion crows (*Corvus corone*) and jungle crows (*Corvus macrorhynchos*) in Kai City and Minami-Alps City, central Japan. Both species were not treated separately because we could not detect differences between them from long distance. The crows were targeted because of the serious damage they cause worldwide (Conover, 1985; McNamara et al., 2002). We employed a fixed 4.8 km line transect method, directing light from a flashlight onto crows within 200 meters of the survey route during daylight hours. Experiments were not conducted in rainy or snowy conditions; surveys were carried out only on clear or cloudy days. The survey time was from 8:30 to 15:00. The survey area was situated at an elevation of 280 meters and was surrounded by rivers, orchards, paddy fields, factories, and residential areas. The survey period extended from November 20, 2023, to March 5, 2024.

### 2.2. Flashlights

In illuminating the crows with a flashlight, we employed multiple flickering patterns that alternated between red and blue light. The flashlight was based on the MAXTOCH OWLEYES W2 Red & Green Beam Version, with the green LED chip replaced by a blue-emitting LED (OSRAM KB CSLNM1.14). Power consumption was 9 watts per LED and the flashlight used two ‘21700’ lithium-ion batteries. The intensity of the light peaked at 450 nm for the blue LED and 624 nm for the red LED. According to the manufacturer’s specifications, the red light has maximum beam distance of 850 meters at night, while there is no available data for blue light.

The flickering frequency was set at three levels: 12.5, 15, and 20 Hz. We used patterns that alternated between red and blue, as well as patterns that included periods of extinguishing (i.e., no light) between the red and blue flickers (Fig. 1). Traditionally, red and blue flickering lights in medical studies do not include a complete off period, always keeping one of the lights on. This method contrasts with flickering monochromatic light, which includes distinct on and off states critical for brain stimulation (Schiller 1992). Our study explores the potential enhancement of the aversive effect by introducing intervals of darkness between the alternating red and blue lights. By adding such off periods (e.g., red light on - all off - blue light on - all off), we investigate whether this modification increases the effectiveness of the stimulus. These dark intervals might more effectively trigger natural brain responses, similar to those activated by the on and off states of monochromatic light. The number of periods of no light per cycle was set at 0-2. Therefore, the illumination patterns consisted of nine variations, combining three flickering frequencies and three patterns of no light. The maximum duration of light exposure was < 60 seconds. If the crows fled, we recorded the time until their escape; if they did not flee, we recorded the maximum exposure time. The behavior of the crows during illumination was recorded with a video camera (HDR-CX370V, Sony, Tokyo, Japan). For each illumination pattern, we conducted experiments over three days. After three days of exposure, we randomly selected the next illumination pattern and repeated the process for all combinations. This procedure was replicated twice, resulting in a total of six days of investigation per pattern.

**Fig. 1.**
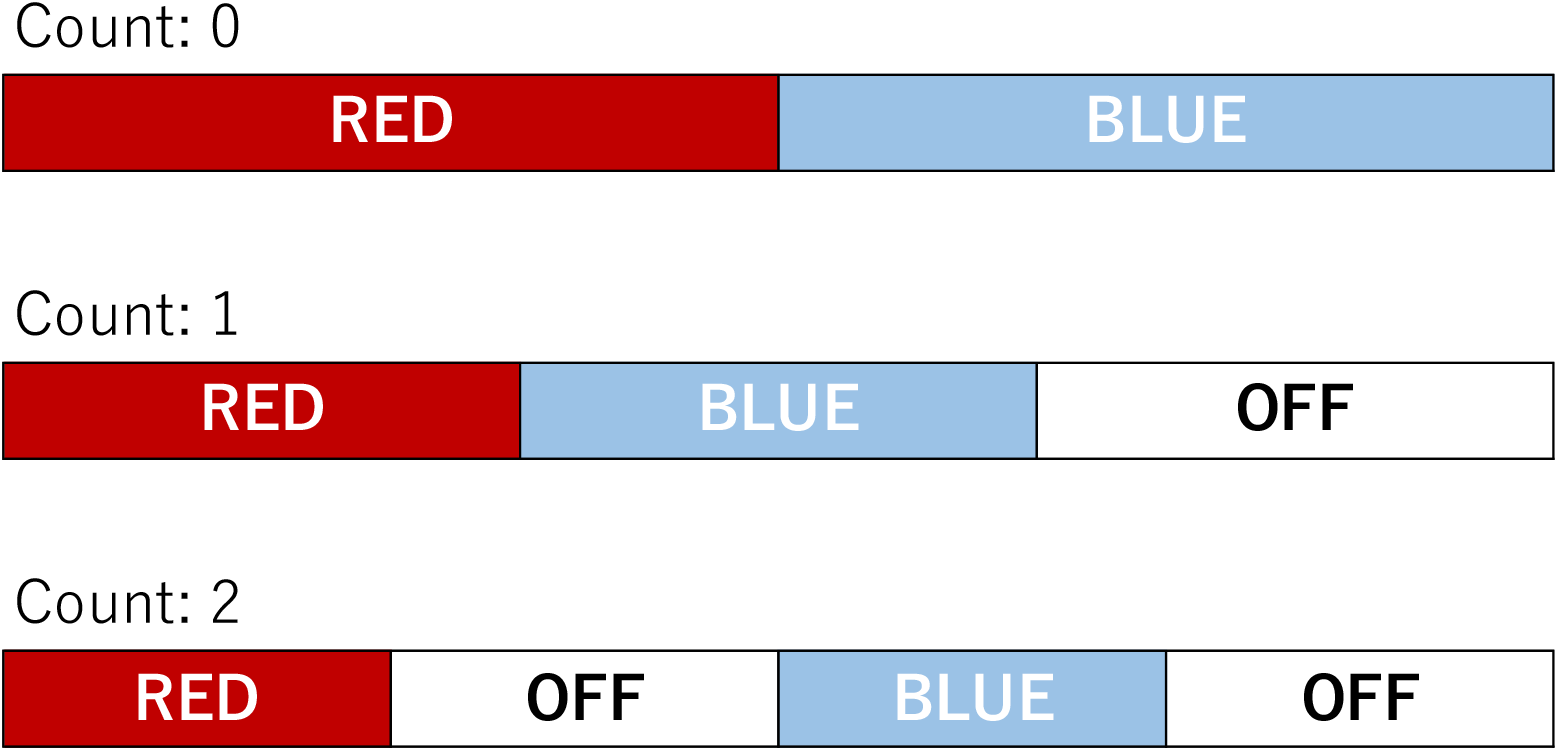
Schematic of the flashing pattern. This figure illustrates the flashing pattern for one cycle, demonstrating that for a frequency of 15 Hz, any of the patterns shown in the upper diagram is repeated 15 times per second. When the number of off counts per cycle is 0, there is no off-time. As the off-count increases, the duration of illumination decreases, even with the same frequency.

### 2.3. Statistical Analysis

The aim of this study was to clarify the effect of different flickering patterns on the escape behavior of birds. For this purpose, we conducted a parametric survival analysis. The dependent variable was the time to escape, and the explanatory variables were flickering frequency, the number of no-light periods, and the distance to the crows. Survival analysis was performed using R software version 4.3.1 (R Core Team, 2023) and the ‘survival’ package. For the regression of a parametric survival model, a log-normal distribution was used.

### 2.4. Ethical notes

This animal study was approved by the Yamanashi Prefectural Research Assessment Committee (#050901).

## 3. Results

After investigating each of the nine flickering patterns, a total of 768 crows were observed. The results of the survival analysis, which examined the time it took for the crows to flee after being exposed to the red and blue flickering light, showed that the model including flickering frequencies and distance was selected over the model based only on distance (likelihood ratio test: Chi-squared = 7.05, d.f. = 3, p = 0.029). The effect of the off-count was not confirmed either from a frequentist perspective (p>0.42) or from predictive values (Fig. 2). The aversion effect was highest at a frequency of 15 Hz, and lowest at 12.5 Hz (Fig. 2, 15Hz vs. 12.5 Hz p=0.011). When illuminated from a distance of 100 meters, the estimated time it took for the crows to flee was 8.1 seconds for the pattern with 15 Hz and two no-light periods.

**Fig. 2.**
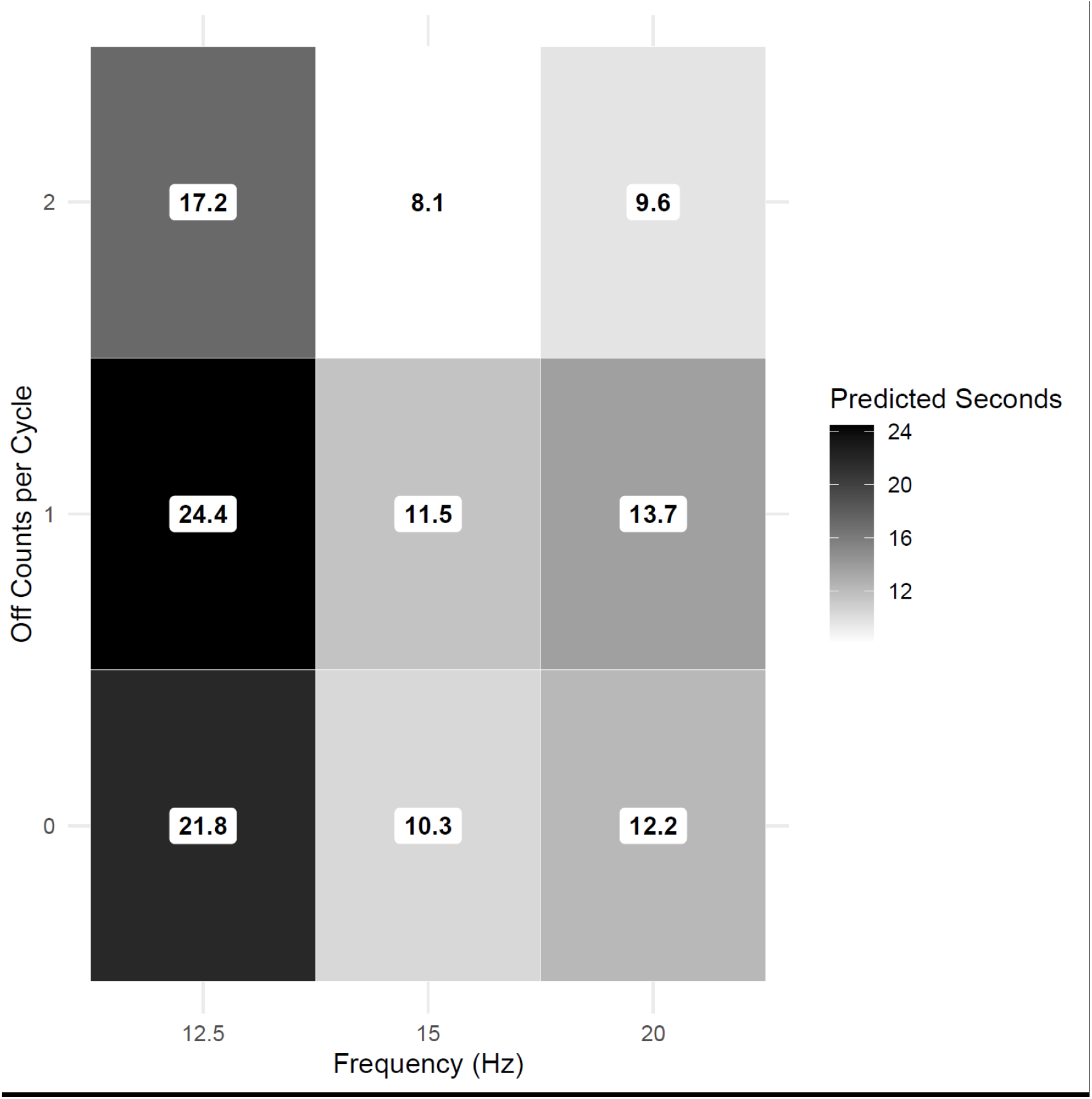
Effects of flashing frequency and pattern on avoidance behavior. This heat plot shows the time taken for avoidance when a flashlight is turned on from a distance of 100m. Darker colors indicate longer times. The numbers within each cell represent the predicted time (seconds).

These results indicate that only flickering frequency affect the escape behavior of crows, and that 15 Hz, which provides the most intense stimulus to the human brain, also provides a similarly strong stimulus to birds. Additionally, the inclusion of a period of extinguishing between the red and blue light phases was found to have no effect on the outcome, suggesting that it is feasible to reduce LED heat generation and power consumption.

## 4. Discussion

The purpose of this study was to assess the patterns of flashing stimuli that are most effective at repelling birds. Our findings suggest that a flashing frequency of 15 Hz, with alternating red and blue lights is the most effective stimulus. Interestingly, the same flashing frequency of 15 Hz is also known to be highly stimulating for the human brain (Parra et al., 2007; Takahashi et al., 1981). It is well-documented that red flashing lights can activate the visual cortex through the red cones, leading to transient abnormal synchronized activity in human brain cells (Tobimatsu et al., 1999; Wilkins et al., 2010). However, there are significant structural differences between the brains of mammals and birds. For instance, the threshold at which a flashing light no longer seems to flicker, known as the critical flicker frequency (CFF), is higher in birds than in humans. (Boström et al., 2016; Potier et al., 2020; Rash, 2004). This indicates that the ability to perceive flashing lights differs between humans and birds, suggesting variations in the neural processing of visual signals from the retina to the brain (Knudsen, 2020; Sharpe and Stockman, 1999). Although there are differences in brain structure, the finding that 15 Hz flashing stimuli affect humans and birds in a similar manner suggests the potential for developing bird-deterrent techniques using visual stimuli, drawing on knowledge from human medical studies.

The alternating red and blue flashing stimuli can be applied in various locations. Not only around agricultural fields but also around airports and wind power facilities, it is possible to repel birds. Additionally, it may be possible to repel birds gathering around ponds near places where poultry are raised, as a measure against avian influenza. Furthermore, its usefulness is expected in preventing wildfires caused by birds perching on power lines. Large birds can sometimes simultaneously contact multiple power lines, and after being electrocuted, their bodies can ignite. This can serve as a source of ignition for wildfires (Barnes et al., 2022). Since power lines are installed almost linearly, illuminating light along the power lines might help reduce house losses due to wildfires.

Our study is not the first report on the interaction between light and bird behaviour. A case similar to the content of this study is one that directly examined the use of flashing lights for bird damage management (Seamans et al., 2001). In their study, a flashing light source with a frequency of 6-8 Hz and mirrors were installed inside the nesting boxes of European starlings. The purpose was to check the deterrent effect (Seamans et al., 2001); however, no decrease in reproductive efficiency due to flashing stimuli was observed, and the potential for light-based damage management was not affirmed. The difference between the results of our study and those of the European starlings is likely due to the significantly different flashing frequencies. In fact, red patrol lights are used in nighttime patrol cars with flashing patterns, yet their impact on the human body is minimal. This implies that the flickering frequency is a crucial factor for both humans and birds.

## 5. Conclusion

Insights from medical and physiological research, we investigated the technology of repelling birds by using flashing light stimuli as aversive conditions. The red-blue alternating stimulus at 15 Hz, known as the most intense stimulus in humans, proved to be equally potent for birds. The flight initiation distance for the crows studied in Japan ranges from 2.6 to 20 meters (Fujioka, 2020; Suzuki and Izawa, 2024). Our finding that avoidance behaviour can be successfully induced within 8 seconds from a distance of 100 m clearly demonstrates the practical effectiveness of this approach in bird management. Moving forward, physiological studies on birds will be necessary to understand how the red-blue alternating stimulus affects the brains of birds. Insights from physiology may reveal the mechanisms that produce aversive stimuli. Additionally, further detailed studies on the interspecies differences in birds’ responses to flashing stimuli will be essential. This study represents the beginning of new technologies that use light as an aversive stimulus. A significant amount of work remains to turn flashing stimuli into practical technology for various purposes and targeting various species.

## CRediT authorship contribution statement

**Takeshi Honda**Writing -review & editing, Writing -original draft, Visualization, Validation, Supervision, Project administration, Formal analysis, Data curation, Conceptualization. **Hiroki Tominaga**: Writing -review & editing, Methodology - Development or design of methodology, Methodology - creation of models. **Akio Shimizu**: Writing -review & editing, Methodology - Development or design of methodology, Methodology - creation of models.

## Acknowledgements

We thank Ms. Kaori Muramatsu for her assistance. During the preparation of this work, the authors utilized GPT-4.0 to improve spelling and grammar.

## Statement

During the preparation of this work the author used GPT-4.0 in order to improve readability and language. After using this tool/service, the author reviewed and edited the content as needed and takes full responsibility for the content of the publication.

